# Transcranial electrical stimulation modulates emotional experience and metabolites in the prefrontal cortex in a donation task

**DOI:** 10.1101/2022.07.31.502214

**Authors:** L Mugnol-Ugarte, T Bortolini, M Mikkelsen, M Carneiro Monteiro, AC Andorinho, Ivanei E. Bramatti, B Melo, S Hoefle, F Meireles, Bo Yao, J Moll, G Pobric

## Abstract

In this study we investigated whether transcranial Direct Current Stimulation (tDCS) applied to two prefrontal cortex regions, the ventromedial prefrontal cortex (vmPFC, anode) and right dorsolateral prefrontal cortex (DLPFC, cathode) can be used to induce changes in self-reported emotions and modulate metabolite concentrations under both electrodes. We employed in vivo quantitative MR Spectroscopy (MRS) in healthy adult participants and quantified the changes in GABA and Glx complex (Glutamate and Glutamine) before and after five sessions of tDCS stimulation. tDCS was delivered at 2mA intensity for 20 minutes for the active group and 1 minute for the sham group. tDCS was applied over five days while participants were engaged in a charitable donation task, known to engage the reward network. We observed increased levels of GABA in vmPFC, but not in DLPFC. Glx levels were decreased in both vmPFC and DLPFC. We found that self-reported happiness increased significantly over time only in the active group. There was a decrease in self-reported guiltiness in both active and sham groups. Altogether, the results indicate that self-reported emotions can be modulated by prefrontal stimulation. This may be because the changes in GABA and Glx concentrations following repeated stimulation induce remote changes in the reward network through interactions with other metabolites, previously thought to be unreachable with noninvasive stimulation techniques.

## Introduction

Every day we encounter complex social environments. Some social contexts are perceived as positive and rewarding, whilst others induce negative feelings such as guilt and regret (Gunaydin & Deisseroth, 2014). Reaction to these contexts can induce a range of behaviors, from prosocial to avoidance to punishing behaviors (Porcelli et al., 2019). One example of prosocial behavior is human cooperation, which have been the focus of behavioral economics, social psychology and, more recently, neuroscience (Fehr & Fischbacher, 2003; Baumeister & Twenge, 2003; Moll et al., 2005; Janowski, Camerer & Rangel, 2013). The pioneering studies of the neural underpinnings of human cooperation have used economic games to establish a basis for investigating human altruistic behaviors (Moll et al., 2006; Harbaugh, Mayr, & Burghart, 2007; Moll et al., 2018, Ruff & Fehr, 2014). Strictly, altruistic behaviors are those voluntarily performed by an agent with the intention to benefit another (non-kin) individual, incurring a cost to the altruistic agent (Trivers, 1971; Zahn, de Oliveira-Souza & Moll, 2020). From an economic perspective these behaviors can be defined as costly actions leading to financial gains for another individual (Fehr & Fischbacher, 2003). Examples of altruistic behaviors depend on the context, and can include blood donations (Zuckerman & Reis, 1978), effort/time spent to help others (Bortolini et al., 2017) or money donations (Eckel & Grossman, 1996).

Indeed, humans often sacrifice material benefits to support social causes (Harbaugh, Mayr & Burghart, 2007), and charitable donations can be used as a proxy for altruistic behavior (Eckel & Grossman, 1996). They can induce the satisfaction derived from voluntary donations (Andreoni, 1990). This feeling of subjective pleasure resulting from altruistic behavior has been related to subcortical areas hypothalamus (George et al., 1995; Moll et al., 2002; Takahashi et al., 2004), prefrontal areas (Funahashi, 2011; Pelletier et al., 2003), and reward and social affiliation circuitry (Moll et al., 2006, Hubbard et al., 2016). This network comprises of dorsomedial and dorsolateral prefrontal cortex (DMPFC and DLPFC), medial frontopolar (mFPC), anterior and subgenual cingulate cortices (ACC and SCC), medial and lateral orbitofrontal cortex (MOFC and LOFC) and medial temporal cortices (Aron et al., 2005; Bickart et al., 2012; Moll et al., 2006). Furthermore, decision making literature has often identified vmPFC (which includes parts of the medial, frontopolar and subgenual PFC) as an area of the brain that is involved in the representation of the value of a stimulus (Arana et al., 2003; Blair et al., 2006; Kable and Glimcher, 2007). Additionally, prosocial decisions recruit subcortical areas implicated in general reward responses, such as the ventral tegmental area (VTA) and the ventral striatum (Inagaki et al., 2016; Moll et al., 2006).

In contrast to prosocial behaviors, failing to help someone or a worthy cause, can lead to feelings of guilt, which has been shown to engage sectors of the vmPFC such as the SCC and mFPC (Wagner et al., 2011). The vmPFC region receives direct cortical connections from the DLPFC. Both vmPFC and DLPFC are connected with subcortical regions involved in emotional responses (Kolb & Whishaw, 2009). Guilt is a powerful emotion that can promote social reparation and prevent socially harmful actions (Moll et al., 2005).

Whilst these fronto-mesolimbic networks play a critical role in prosocial behaviors, the combined role of the specific neurotransmitters that mediate these functions is still not well understood (Ongur and Price, 2000; Depue Morrone-Strupinsky, 2005). Animal studies show a critical role of dopamine (DA) in prosocial behaviors (Gunaydin & Deisseroth, 2014). DA in the VTA increases its activity in response to social stimuli, and the degree of DA release is associated with the duration of social interaction (Scott-Van Zeeland et al., 2010). Moreover, the gamma-aminobutyric acid (GABA; Karreman & Moghaddam, 1996) and glutamine (Gln) concentrations in the prefrontal cortex have an important role in modulating activity and DA release in the midbrain and striatum (Carr & Sesack, 2000). Additionally, oxytocin (OXT) regulates social reward, by enhancing VTA DA signals through GABA system stimulation (Eisenberger, 2012).

Non-invasive brain stimulation techniques such as transcranial Direct Current Stimulation (tDCS) (Pascual-Leone, Walsh & Rothwell, 2000) are increasingly used to study causal involvement of brain areas. tDCS modulates cortical excitability in the underlying cortex by “up-regulating” or “down-regulating” a region of interest (Nelson et al., 2014). At currents of 1 mA, generally, tDCS results in depolarisation of the neurons underneath the anode, hence causing an excitatory effect. In contrast, tDCS causes hyperpolarization underneath the cathode and thus inhibition of cortical neurons in the motor cortex (Nitsche & Paulus, 2000). Using 1mA protocols, tDCS has been shown to modulate emotional pain (Boggio, Zaghi and Fregni, 2009), negative emotion perception and to boost emotion regulation (Vergallito et al., 2018). Importantly, tDCS applied to the MPFC was demonstrated to influence feelings of guilt and change participant’s willingness to perpetrate social violations (Karim et al., 2010).

Applying tDCS at current strength of 2 mA, however, causes excitability increases under both anode and cathode (Batsikadze et al., 2013). It is less clear whether the reversal of inhibitory effects following 2 mA stimulation holds for other cortical regions such as the prefrontal cortex. Magnetic Resonance Spectroscopy (MRS) can be combined with tDCS in order to study changes in metabolite concentrations following stimulation. Major excitatory and inhibitory neurotransmitters Glu and GABA have been reported to be involved in tDCS secondary effects (Fritsch et al., 2010; Stagg et al., 2009). For instance, GABA is involved in the anodal tDCS after-effects, while both GABA and Glu concentrations have been modulated following cathodal stimulation (Polanía, Nitsche & Ruff, 2018).

Notably, there are few studies that applied prefrontal tDCS and measured metabolites mostly under the anode (Mezger et al., 2021). For example, anodal tDCS at 1mA has decreased the Glx levels (Glutamate and Glutamine) in the DLPFC (Guan et al., 2020) and anodal tDCS at 1mA decreased GABA concentration in the premotor cortex (Bachtiar, 2015). Other studies report no effect of stimulation on Glx quantification (e.g. Hone-Blanchet, Edden & Fecteau, 2016). Numerous studies have shown a facilitatory effect of anodal tDCS of 2 mA on motor learning and skill (Galea et al., 2009; Grimaldi et al., 2014; Cantarero et al., 2015). Therefore, it is still not clear the effects of tDCS on the cognitive process of monetary donation to charity.

In this study, we used tDCS in combination with MRS to explore the neural mechanisms of two emotions associated with altruistic donations, by modulating neural activity in vmPFC. We used a specific tDCS electrode montage where both electrodes were stimulating (Chib et al., 2013), which has been shown to modulate midbrain regions following stimulation of the PFC. tDCS was applied at 2mA during a modified Dictator Game, in which participants decided how much money they would like to donate to different Non-Governmental Organizations (similar to Moll et al., 2006). First, we tested whether tDCS would have an effect on happiness and guiltiness in the context of costly altruistic decisions. Stimulation was expected to increase perceived happiness, while diminishing perceived guilt. Importantly, MRS was used to estimate the effect of 2mA stimulation on metabolite concentration in vmPFC over both anode and cathode. We expected that, over five sessions of non-invasive brain stimulation, changes will be observed in brain metabolites (GABA and Glx) in vmPFC and DLPFC, where electrodes were positioned. In line with Dickler et al., (2018) we expected increase of GABA following 2mA tDCS stimulation over vmPFC and decrease of Glx in the DLPFC (Guan et al., 2020).

## Methods

### Participants

41 Participants (20 women) living in Rio de Janeiro, Brazil (Mean age = 24.4 ± 10; range = 19 to 34 years-old) took part in the study. Due to the constraints of the neurostimulation and MRI protocols, personal history of epilepsy, cardiac pacemaker, previous intracranial surgery, pregnancy, regular psychotropics intake and inability to give informed consent were the exclusion criteria. Because of the complexity of the tasks, the educational level was used as an inclusion criteria: participants were undergraduate students or held a university degree. The study was approved under ethics protocol number 2.036.768 at D’Or Institute for Research and Education, Rio de Janeiro, Brazil, where the research was conducted.

### Protocol overview

A between-subjects design was employed, in which participants took part in a donation task while one group received tDCS and the other group received sham stimulation on 5 consecutive days. Glx and GABA spectroscopy, functional Resting State rs-fcMRI and Diffusion Weighted Image (DWI) were acquired on days 1 and 5 (rsfc-MRI and DWI results will be reported elsewhere). Experimental groups were pseudo-randomly formed as follows. The first participant received an identification number and was randomly allocated to one of the experimental groups, the next participant was allocated to the other group and so on, always pairing the sex ratio in both groups. The task was performed concurrently with the tDCS for five days (Fig. 1).

**Figure 1:**
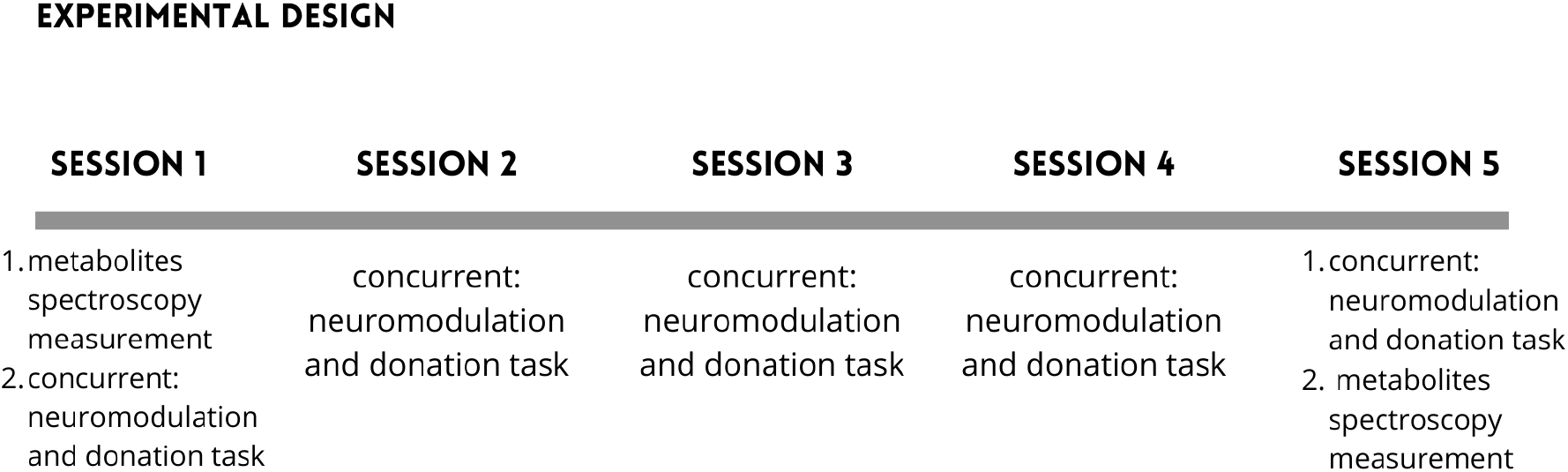
Experimental design overview

### MRS Acquisition

Magnetic Resonance (MR) images and spectra were acquired on a 3T PRISMA scanner (Siemens Healthcare, Erlangen), using a 64-channel receive-only head coil. After the recording of a scout image, high-resolution anatomical images were acquired using a three-dimensional T1-weighted magnetization-prepared rapid acquisition gradient echo MPRAGE sequence (repetition time (TR), 1800 ms; echo time (TE), 2,26 ms; inversion time, 900 ms; flip angle, 8 deg; 256 × 256 matrix; 1 mm^3^ isotropic voxel; 176 slices in sagittal orientation with no gap; FOV 256 mm). MRS images were acquired on Day 1 before the tDCS and donation task and on Session 5 after the tDCS and the task. More information about Spectroscopy is at Supplementary Material 2.

### Transcranial Direct Current Stimulation (tDCS)

Participants were either stimulated with tDCS (Active group) or they received sham stimulation (Sham group), while they took part in a donation task (details below). DC-STIMULATOR PLUS (NeuroConn, GmbH) electrodes were placed on the corresponding area 10 (anode) and right lateral BA 9, which comprehends dorsolateral prefrontal cortex (rDLPFC; cathode, Fig 2); Following Chib et al., (2013) procedure, the electrodes measured 20 cm^2^ and 15cm^2^, the sponges were wet with saline solution and the current intensity was 2mA; Both groups were instructed that they would be stimulated for 20 minutes while they performed a decision-making task. Firstly, participants, would start the task, after 10 minutes, the Active group received a stimulation of 2 mA during 20 minutes with a ramp up and a ramp down of 30 seconds, while the Sham group was stimulated similarly as in Antonenko et al., (2017) with a ramp up of 30 seconds, stimulation of 1 minute and a ramp down of 30 seconds. In order to confirm topographical effects of neuromodulation, we modeled the magnitude of the total electric field due to stimulation with ROAST (Jung et al., 2013). The model provided evidence that the tDCS electric field was largest over the right vmPFC region.

**Figure 2:**
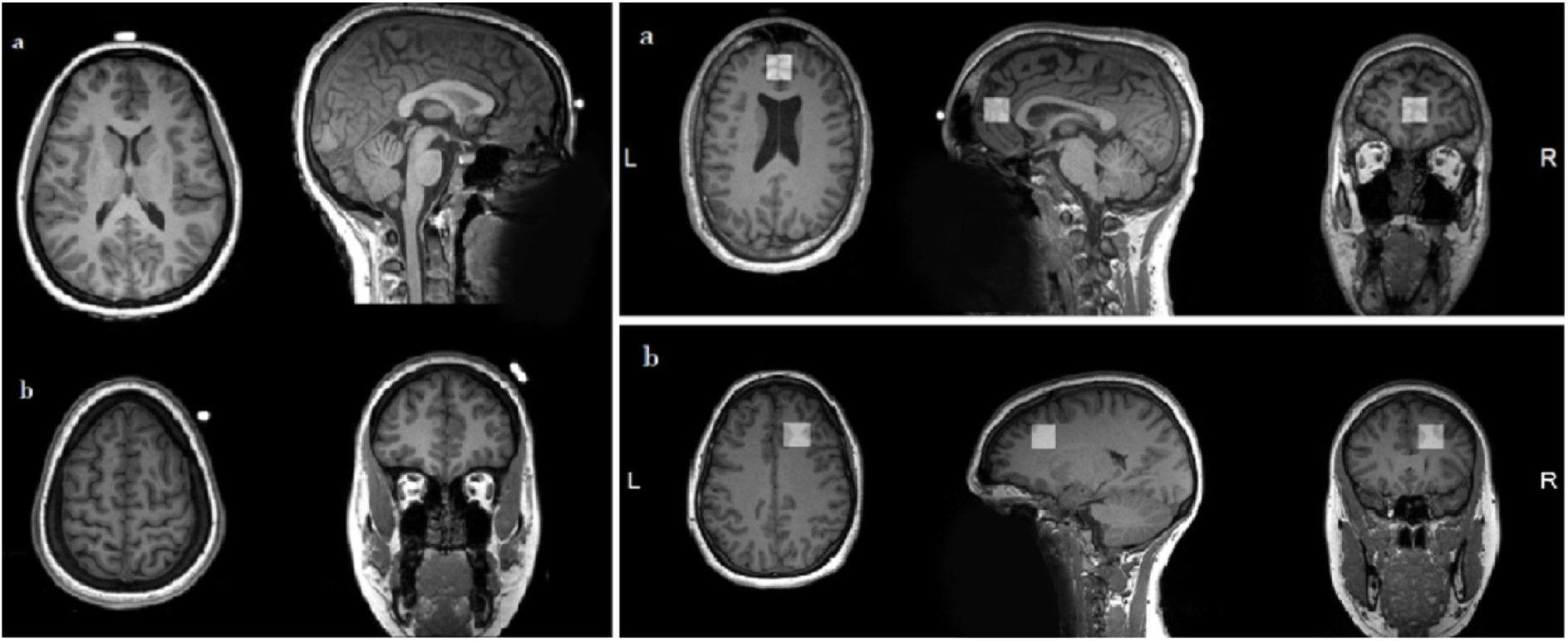
example of MRS voxel positioning on (a) medial BA 10 and (b) lateral BA 9 from one participant.

### Donation task

The task was delivered in Presentation® (Version 18.0, Neurobehavioral Systems, Inc., Berkeley, CA, www.neurobs.com). In a modified Dictator Game, 50 Brazilian Non Governmental Organizations (NGOs) were presented per day throughout 5 days of experiment, totaling 250 NGOs. The task consisted of the participants deciding how much money they wanted to give to each NGOs. A hundred NGOs were real, while the remaining were created solely for experimental purposes. The created NGOs were described in a similar way as the real ones (see Figure 4) and participants were informed that all NGOs were real. The NGOs supported different causes, with similar proportion of types of NGO: animals welfare 29.2%), humanitarian (53.6%) and controversial (17.2%; i.e.: pro guns, ethnic equality, abortion, etc.). Before starting the experiment, the participants read an explanation sheet about the task (Supplementary material; Text 1) and after confirming verbally that they understood the task, the tDCS electrodes were positioned on their heads. They were informed that they had earned R$50 and could donate any amount from $0 to $50 to each of the 50 NGOs presented to them per day along the 5 days. In addition, they were informed that on the last day one of the 250 donation trials would be drawn and the amount given on that trial would be donated to the respective NGO, and the remaining amount would be given to the participant.

**Figure 3:**
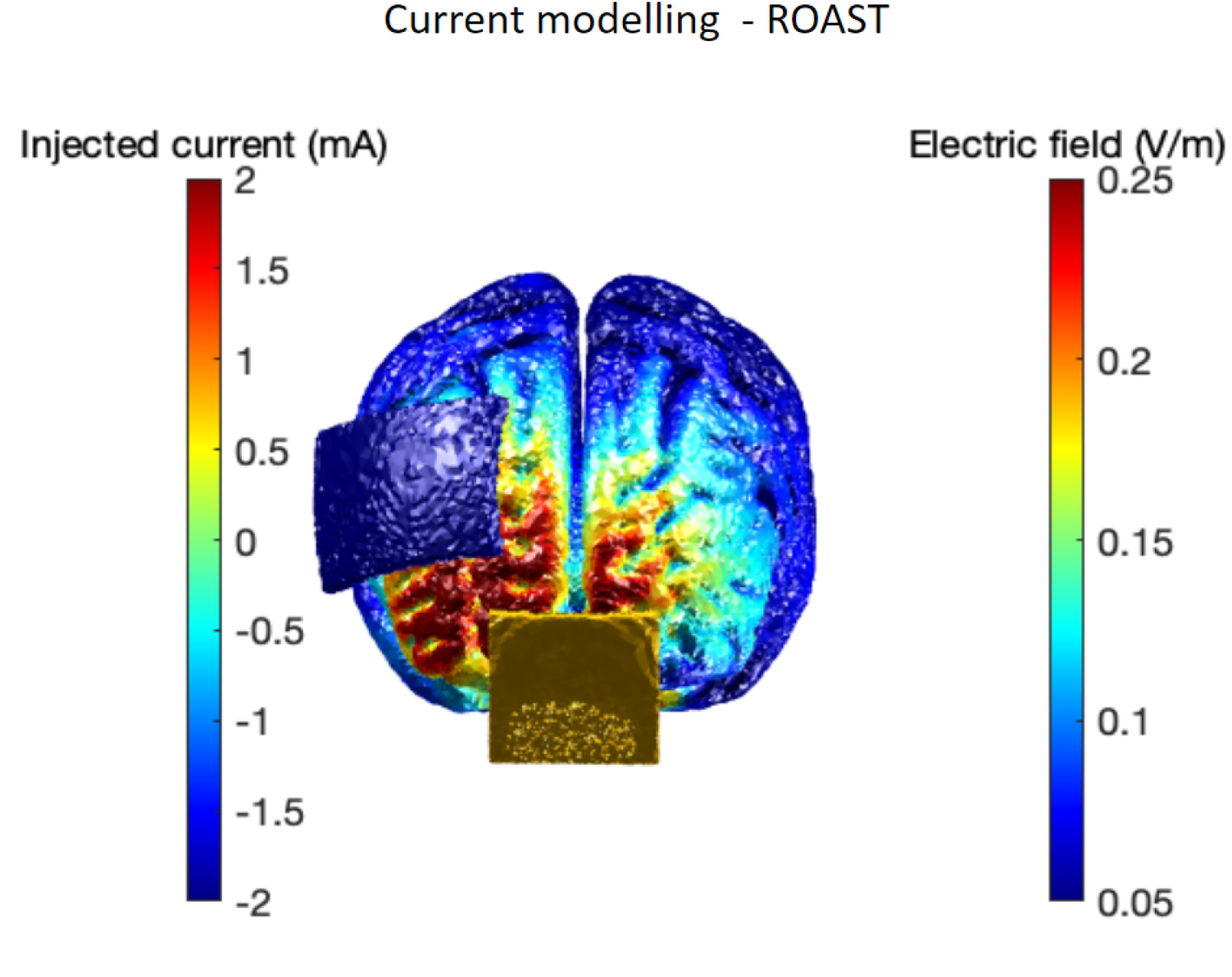
model of the magnitude of the total electric field due to stimulation was made with ROAST (Jung et al., 2013).

**Figure 4:**
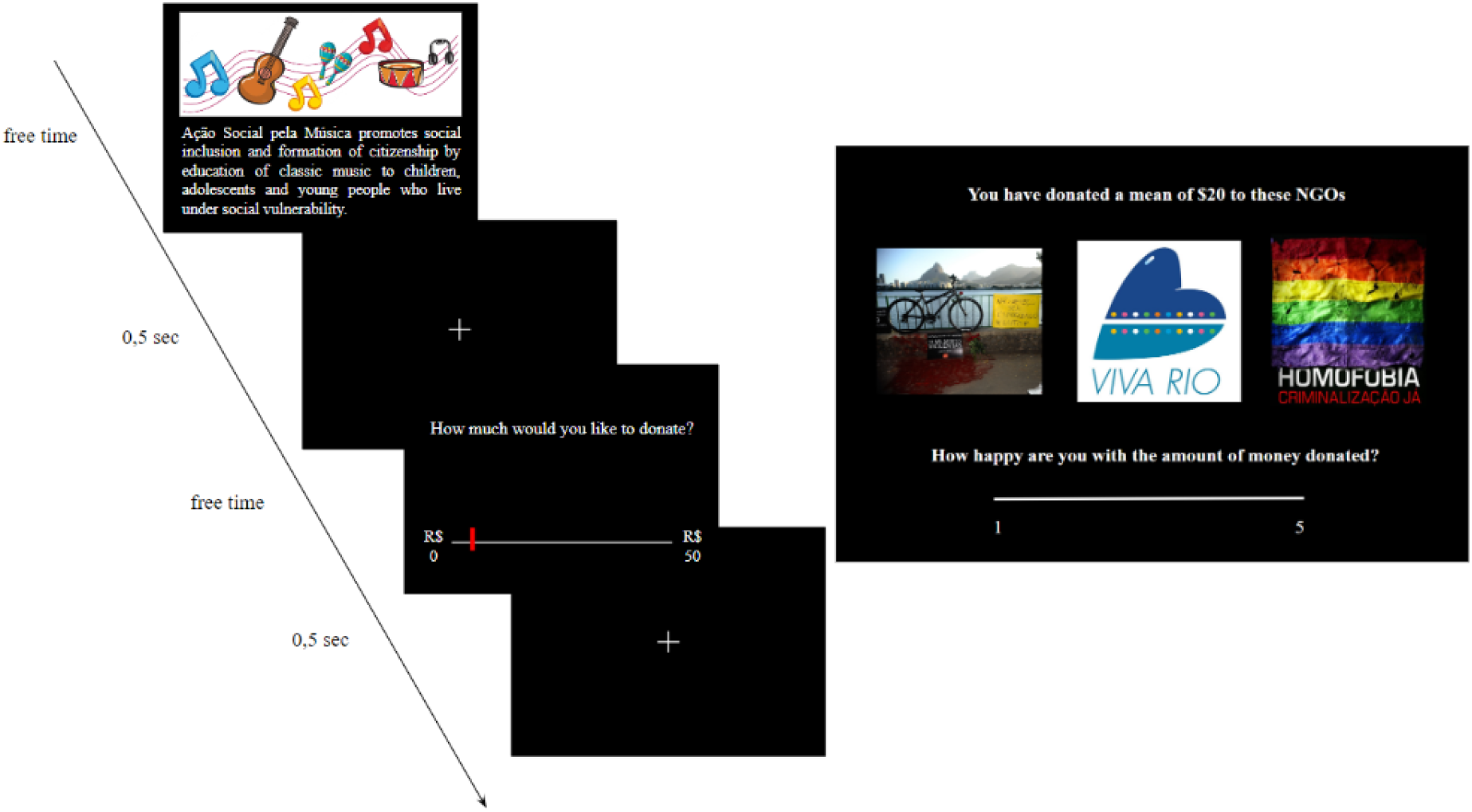
A) Donation task - On the first day there were 2 training trials with the same structure as the actual task. In each trial participants were presented with a photo representing the NGO, its name, and a brief description of its cause and target public audience (e.g. “*Ação Social pela Música* promotes social inclusion and formation of citizenship by education of classic music to children, adolescents and young people who live under social vulnerability”. After reading the description with no time constraints, participants had to decide whether to donate any amount of money ranging from 0 to 50 Brazilian Reais for each NGO. B) Happiness and Guiltiness questions; at the end of the donation task, participants were asked about how happy and guilty they felt after choosing to donate specific amounts of money to a variety of different NGOs.

At the end of the whole task participants self-reported how happy and guilty they felt (from 1 to 5) regarding the amount of money donated for different NGOs. The NGOs were grouped in quartiles of participant’s mean donations. Therefore, the NGOs that received 25% of the total amount of money donated by the participant represented the 1st quartile of money donated; the 2nd quartile was the range representing 25 to 50% of the total amount of money donated, and so on).

### MR Spectroscopy Analyses

Edited spectra were analyzed using Gannet (**Edden et al., 2014**) with a measured basis set containing GABA, Glx complex (Glutamate and Glutamine) and Creatinine (Cr); no water reference data was collected. Nonlinear least-squares fitting was used to model the difference spectrum between 2.79 and 4.10 ppm with a three-Gaussian function with a nonlinear baseline to quantify the 3.0 ppm GABA signal, and the 3.75 ppm Glx complex signals (**Mikkelsen et al., 2017**). Quantification of GABA was estimated as the integral ratio between GABA+ and Cr. This ratio (hereafter “GABA”) was used as the variable of interest in the analysis described below (**Edden et al., 2014**).

Each MRS dataset was visually inspected for data quality and signal artifacts. Data exclusion criteria was based on **Waddell et al., 2007**. A total of 18 datasets (11 women; 8 from the active group) were excluded based on poor SNR of the 3 ppm GABA+ peak or high model fit error (>20%) from vmPFC and 4 datasets (1 woman from Sham group; 3 men: 2 from Sham group, 1 from Stim group) with poor signal from DLPFC; 3 excluded datasets were in common. Because of these exclusions, all the analyses - MRS and behavioral - were run with the participants that have good spectroscopy signals. The higher proportion of excluded participants based on the vmPFC signal is expected given the higher susceptibility effects in this region, which affects signal to noise ratios. More info can be accessed in Supplementary Material 4.

## Results

### 1. The effects of Session and Stimulation on reported happiness and guiltiness

To illustrate how self-reported happiness and guiltiness changed over time, we plot the mean happiness and guiltiness ratings as a function of time in and Active group in Figure 5.

**Figure 5.**
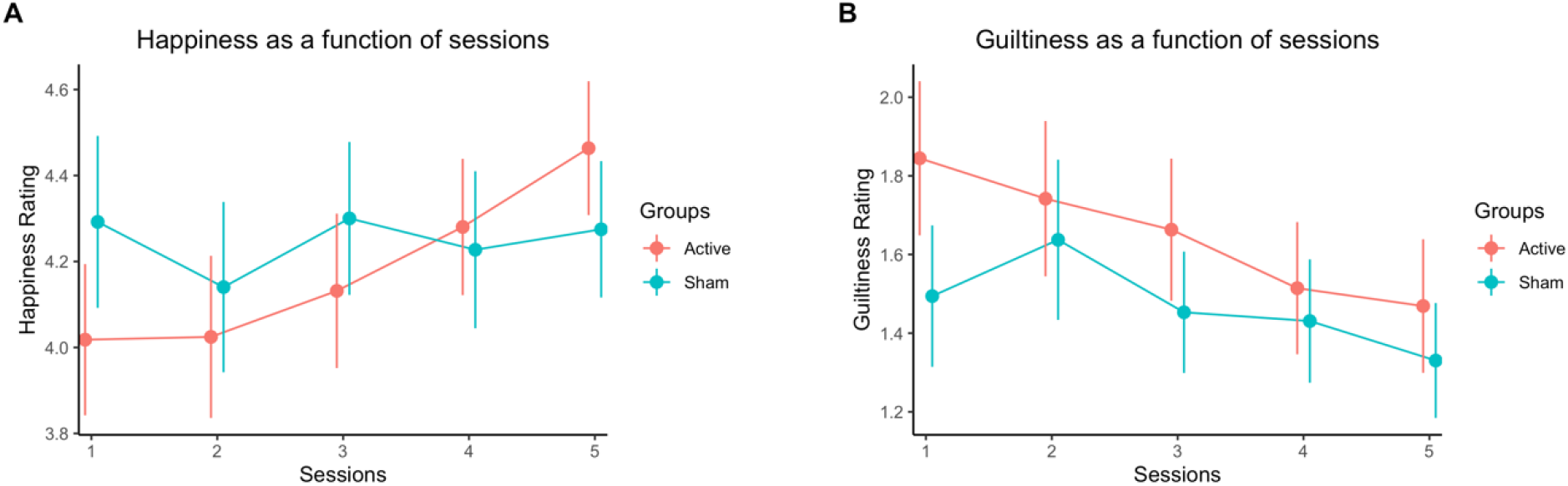
Mean happiness and guiltiness ratings over time and between groups; error bars indicate standard errors.

To assess the effects of session and stimulation on reported happiness and guiltiness, we fitted linear mixed-effects models (LMEMs) for each respective measure using the lmer() function of the lme4 package (Bates et al., 2015). Both models shared the same fixed-effect and random-effect structures. The former included fixed factors of Time (1,2,3,4,5 days) and Group (active, sham) and their interactions; both factors were deviation coded, with a mean of 0 and a SD of 0.5. The latter employed the maximal random-effect structure by design, including a by-Subject random intercept and a by-Subject random slope for Time. The *p*-values for fixed effects were computed using Satterthwaites’s approximation using the lmerTest package (Kuznetsova, Brockhoff, & Christensen, 2017).

For reported Happiness, there was a significant main effect of Session (*b*=.178, *SE*=.064 *t*=2.773, *p*=.009), and a significant Session × Group interaction (*b*=.311, *SE*=.129, *t*=2.417, *p*=.021). Specifically, reported happiness increased significantly over time, however only in the Active group (*b*=.325, *95%CI*=[.146 .504]) but not in the Sham group (*b*=.015, *95%CI*=[-.173 .203]). The main effect of Group was not significant (*p*=.712), suggesting that happiness did not significantly differ between groups overall which may be explained by lower starting happiness in the Active group.

For reported Guiltiness, there was a main effect of Session (*b*=-.218, *SE*=.080, *t*=-.2.735, *p*=.009). The main effect of Group and the Session × Group interaction were not significant, *p*s>.287. These suggest that reported guiltiness decreased significantly over time, independent of stimulation.

### 2. MRS analyses: the effects of intervention and stimulation on GABA and Glx levels at vmPFC and DLPFC

To illustrate the effects of intervention, we plot the changes in GABA and Glx levels post intervention (session 5) relative to the baseline (session 1) by Group (active, sham) and Region (vmPFC, DLPFC) in Figure 6.

**Figure 6.**
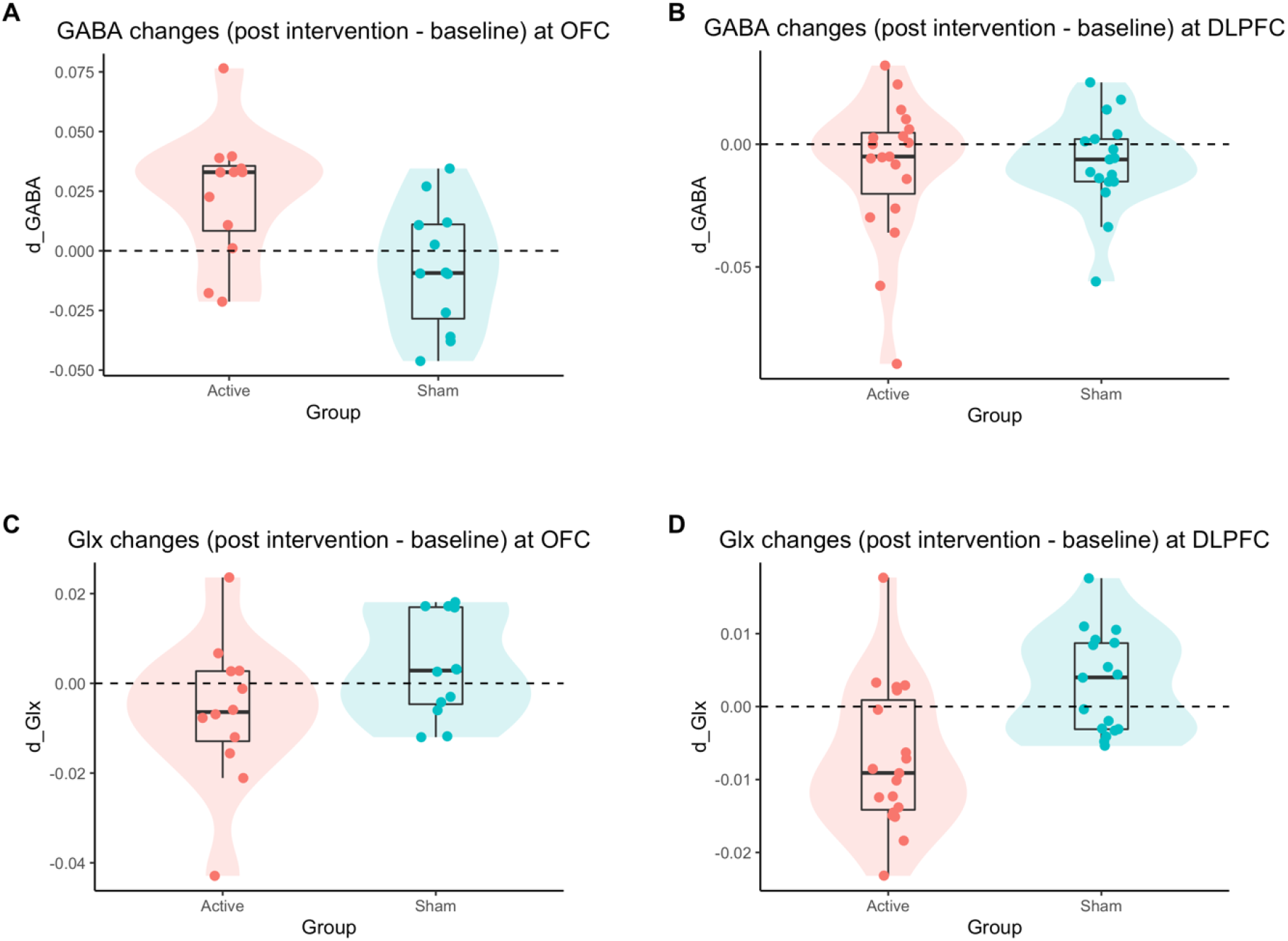
Changes of GABA (upper row) and Glx (lower row) post-intervention in Active group at vmPFC (left column) and at DLPFC (right column). Jittered dots represent individual observations in each group, with their distributions illustrated by the underlying violin shapes. Boxplots indicate the median and the 1^st^ and 3^rd^ quartiles of the distributions.

To assess the effects of intervention, Active group and stimulation site on GABA and Glx levels, we fitted separate linear mixed-effects models (LMEMs) for each respective neurotransmitter using the lmer() function of the lme4 package (Bates et al., 2015). Both models shared the same fixed-effect and random-effect structures. The former included fixed factors of Intervention (post-intervention - baseline), Group (stimulation, sham) and Region (vmPFC, DLPFC), as well as their interactions; all factors were deviation coded, with a mean of 0 and a SD of 0.5. The latter employed the maximal random-effect structure by design, including a by-Subject random intercept and a by-Subject random slope for Intervention. The *p*-values for fixed effects were computed using Satterthwaites’s approximation using the lmerTest package (Kuznetsova, Brockhoff, & Christensen, 2017).

For GABA, there was a significant two-way interaction between Intervention and Region (*b*=1.730e-02, *SE*=6.669e-03, *t*=2.594, *p*=.011). While GABA levels decreased significantly post intervention at DLPFC (*b*=-.009, *95%CI*=[-.017 -.000]), they did not significantly respond to intervention at vmPFC overall (*b*=.008, *95%CI*=[-.002 .019]). The lack of intervention effects at vmPFC could be explained by a significant three-way interaction between Intervention, Group and Region (*b*=3.323e-02, *SE*=1.334e-02, *t*=2.491, *p*=.014): at vmPFC, the effects of Intervention significantly varied by Group (*b*=.031, *SE*=.010, *t*=2.997, *p*=.003), as GABA levels significantly increased in the Active group (*b*=.024, *95%CI*=[.009 .038]) but not in the Sham group (*b*=-.007, *95%CI*=[-.022 .008]); At DLPFC, although GABA levels significantly decreased overall (*b*=-.009, *SE*=.004, *t*=-2.039, *p*=.044), but the effects of intervention did not interact with Group (*p*=.788), indicating that the decreases in GABA were not driven by tDCS. Taken together, the results suggest that GABA levels significantly increased in response to tDCS to vmPFC, but not to DLPFC.

For Glx, there was a significant main effect of Region, showing that Glx levels were overall higher at vmPFC than at DLPFC (*b*=.012, *SE*=.002, *t*=7.209, *p*<.001). There is an Intervention x Group interaction (*b*=-.010, *SE*=.003, *t*=-3.271, *p*=.002), indicating that Glx levels significantly decreased in the Active group (*b*=-.007, *95%CI*=[-.011 -.002]) but did not significantly change in the Sham group (*b*=.003, *95%CI*=[-.001 .008]). The results suggest that Glx levels significantly decreased after tDCS to both vmPFC and DLPFC.

## Discussion

In this study we used tDCS over both vmPFC and DLPFC, to examine changes in GABA and Glx (Glutamate and Glutamine) while participants engaged in a donation task. We have shown that the simultaneous tDCS anodal stimulation over the vmPFC and cathodal stimulation over the DLPFC influenced the self-reported emotions following a donation-task. There was an increase of happiness and a tendency of decrease of guiltiness in the Active group across sessions. Importantly, we report increased levels of GABA comparing initial and final spectroscopic levels of this metabolite in the vmPFC as a result of 2 mA tDCS stimulation over the week on the Active group.

The Glx level was decreased in both vmPFC and DLPFC. The observation that the repeated tDCS causes the decrease of excitatory neurotransmitters implies the different underlying mechanisms between the repeated tDCS and the single stimulation, in line with Guan et al. (2020). Furthermore, cathodal stimulation of DLPFC might have suppressed its control over vmPFC, which resulted in the enhancement of the anodal effect of the latter. This enhanced vmPFC stimulation might have yielded an increased remote activation of the distally interconnected ventral midbrain, which in turn could disinhibit subcortical dopamine release. Both direct GABA decrease and indirect Glx remote activation most likely manifested behaviorally as increases in participants’ happiness ratings.

The increase of happiness of the Active group across sessions, and a tendency for a decrease of guilt suggests that the tDCS anodal stimulation over the vmPFC and DLPFC modulates the experience of these two emotions. This interpretation is in line with the notion that the vmPFC – which includes the MOFC (Nejati et al., 2021) is crucial for emotional experience in social contexts. In fact, the MOFC plays a pivotal role in emotion in relation to reward values, integrating sensory and abstract aspects of stimuli to behavioral goals (Rolls, 2019). The MOFC also encodes emotional stimuli (Winecoff et al., 2013) and is part of the emotional immediate choices processing (Manuel, Murray & Piguet, 2019).

Due to the size of the electrodes and the bipolar scalp electrode organization, there are intrinsic spatial uncertainties in measuring the effects of tDCS on emotions and behavior (Nitsche et al., 2008). In this study the tDCS stimulation evoked changes only in reported happiness of the Active group, even though tDCS applied to the MPFC was demonstrated to influence feelings of guilt (Karim et al., 2010). It is well established that tDCS does not only affect the brain regions directly under the electrodes but may also modulate connectivity among remote and functionally associated brain areas (Polanía, Nitsche, & Paulus, 2011) by influencing the strength of network connectivity (e.g. Meinzer et al., 2012). Furthermore, since the MPFC is associated with representing other’s beliefs and emotions and related to social cognition processes (Takahashi et al., 2004), the donation task could have induced the rise of self-perception of happiness.

We reported a tendency of decrease of guilt experience in the Active group over time. Feelings of guilt are associated with activation of the MPFC, among other brain regions (Takahashi et al., 2004). Furthermore, the FPC (BA 10) - which is part of vmPFC (Kolb & Whishaw, 2009) has been consistently involved in moral judgments (Moll et al., 2001; Heekeren et al., 2005) and prosocial sentiments such as guilt (Moll et al., 2002; Moll et al., 2007; Zahn et al., 2009c). Therefore, the tDCS anodal stimulation could be influencing the feeling of guilt directly at the prefrontal cortex or through its interconnection with other frontal and subcortical circuits.

One caveat worth mentioning is that participants were required to remain immobile in the MRI scanner for up to 90 minutes, as the slightest movements could negatively affect the MRS signal negatively; this is especially critical for MRS obtained from the prefrontal cortex next to brain-bone-air interfaces, which was the case in our study. Strict analysis of signal distortion led to loss of almost 50% of the spectroscopy data, which further reduced the power of our study in regard to the MRS findings. Considering that the neuroanatomy and scalp electrode impedance (tissue resistance using electrodes) of each participant are different, another caveat to be mentioned is the difficulty in controlling the current flow in each specific stimulated region affected by the tDCS electrodes. Both prefrontal regions cannot be precisely located and stimulated, as they are in places conducive to signal interference.

Also, the effects on Glx should be interpreted with caution, as the Glx complex describes the contributions of two metabolites, i.e. Glutamate and Glutamine wherein glutamate concentration in the brain is up to 45% higher than Glutamine concentration (Jang et al., 2005). Moreover, glutamate is not only the primary excitatory neurotransmitter in the brain, but it is also implicated in the amino acid synthesis of GABA (Michels et al., 2012; Rae, 2014). Thus, given that the MRS Glx signal encloses contributions from several glutamate pools, it was not possible to separate the spectral contributions resulting from the neurotransmitter population of Glutamate propper from those resulting from the other Glutamate pools.

Our findings imply that anodal tDCS of vmPFC and cathodal stimulation of DLPFC can be used to induce remote changes in regions deep within the brain, which were conventionally thought to be unreachable with noninvasive stimulation techniques.

Decision-making tasks (similar to our donation task) that require higher level of reasoning often recruit DLPFC (Krain et al., 2006) which was the location of our cathodal electrode. With this in mind, future work must take into account how more complicated behavioral tasks might interact with electrode placement and polarity. In conclusion, we provide a test-case of how a network of interconnected prefrontal brain areas can be stimulated with tDCS to promote prosocial behavior and/or change emotional responses associated with prosocial decisions. Our findings imply that anodal tDCS of vmPFC and cathodal stimulation of right DLPFC can be used to induce remote changes in the reward network through interactions with other metabolites in regions deep within the brain, which were conventionally thought to be unreachable with noninvasive stimulation techniques.

## Supplementary Material

### 1 Donation task explanation

**Hello!**

Thank you very much for your participation, science thanks you.

You will participate in a task where you will donate money to NGOs. In each donation you will have R$50 available to donate. We ask that you treat each donation **as unique**.

Make each donation as if you were using your **own money**. Please use the entire donation scale (from R$0 to R$50), that is, evaluate how much each NGO **deserves**.

At the end of the experiment you’ll see the summary of your donations. At this point we will ask you how happy or guilty you felt when making those donations. Please be aware of these two feelings and indicate your response on the scale.

On the last day of the experiment we will randomly assign 1 NGO that we will donate the money corresponding to your choice, and you will get the rest.

In addition, by participating, you will receive R$100

- R$12 per day (x 5) for participation = R$60
- R$8 per day (x 5) for the transportation to the institute = R$40.

### 2 Spectroscopy Acquisition

This image was used to place the voxel of interest (20 × 20 × 20 mm^3^) over the corresponding areas of the vmPFC and the DLPFC (**Yousry et al., 1997**) based on capsules placed over the two corresponding regions of the scalp (Fig. 2). The medial voxel was positioned in the parenchyma in front of the genu of the corpus callosum, aligned with the vmPFC. The other voxel was placed in the region comprehending the dorsolateral prefrontal cortex (DLPFC), part of the dorsolateral prefrontal cortex. For MR Spectroscopy (MRS), first the transmitter radio frequency voltage was calibrated for the individual volume of interest, followed by the adjustment of all first - and second-order shims using FAST(EST)MAP (**Gruetter, 1993; Gruetter & Tkác, 2000**). GABA-edited spectra were recorded using the MEGA-PRESS technique (**Mullins et al., 2014**). 80 scans were acquired with a TR of 2000 ms and a TE of 68 ms. Immediately afterwards, a spectrum without water suppression was recorded (eight scans).

### 3 Transcranial Direct Current Stimulation (tDCS)

The impedance was recorded for each participant, and the mean impedance of the subgroup of analysis was ∼3,17. The electrodes position was defined and estimated by the program Beam F3 (**Beam et al., 2009**) which takes into account three head measures: circumference ear to ear, over top and inion-nasion.

### 4 Metabolites Analysis

The analysis consisted of four steps: 1) GannetLoad, in which the metabolites were provided including OFF and ON scans; 2) GannetFit, which estimated the area under the edited GABA signal at 3 ppm and Cr signal at 3 ppm as shown in Supplementary material 5; 3) GannetCoRegister, which registered the MRS voxels to the T1-weighted image; and 4) GannetSegment, which segmented the T1-weighted image into gray matter, white matter and cerebrospinal fluid (CSF) for the voxel and a CSF-corrected GABA estimate.

A sample magnetic resonance spectrum for GannetFit

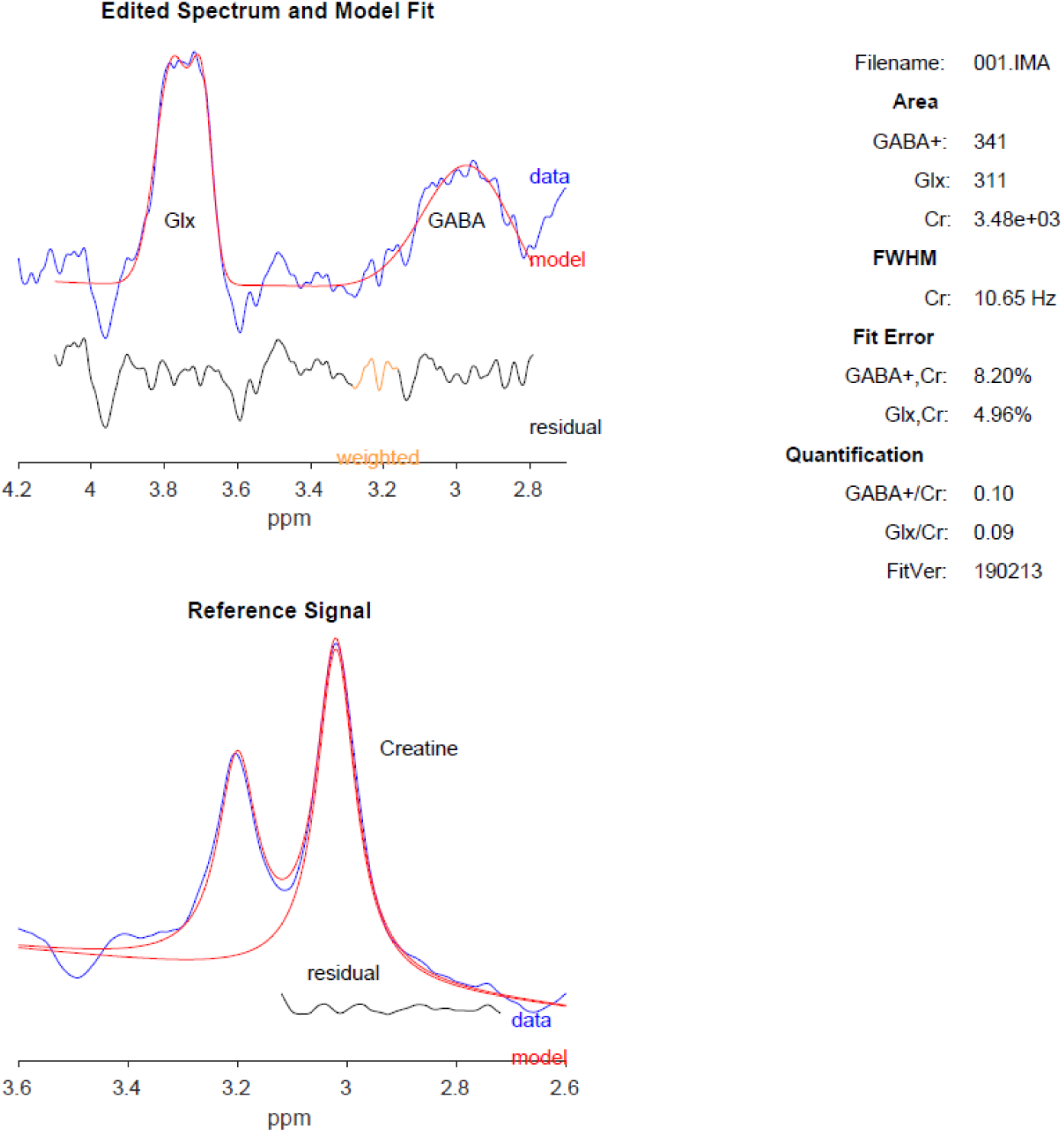

