## Supplementary Material for "Transcranial electrical stimulation modulates emotional experience and metabolites in the prefrontal cortex in a donation task"

A sample magnetic resonance spectrum for GannetFit

**Edited Spectrum and Model Fit**

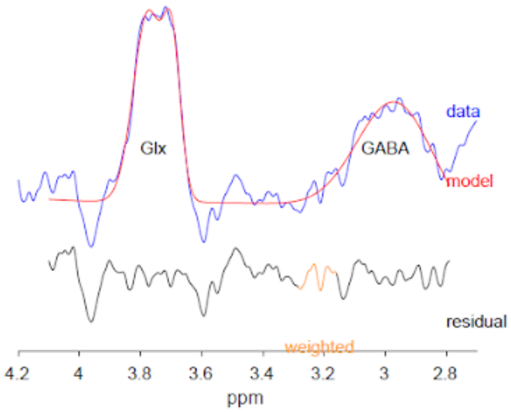


| | |
|---|---|
| Filename: | 001.IMA |
| **Area** | |
| GABA+: | 341 |
| Glx: | 311 |
| Cr: | 3.48e+03 |
| **FWHM** | |
| Cr: | 10.65 Hz |
| **Fit Error** | |
| GABA+,Cr: | 8.20% |
| Glx,Cr: | 4.96% |
| **Quantification** | |
| GABA+/Cr: | 0.10 |
| Glx/Cr: | 0.09 |
| FitVer: | 190213 |

**Reference Signal**

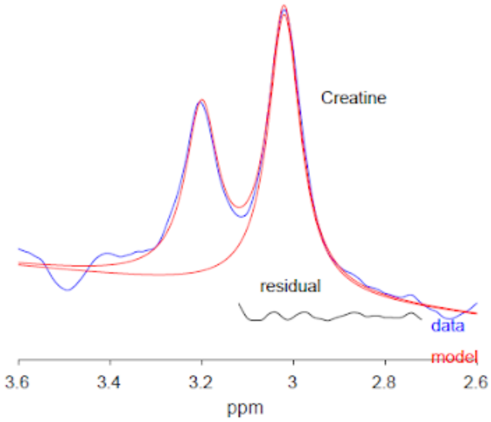
